# Penguin: A Tool for Predicting Pseudouridine Sites in Direct RNA Nanopore Sequencing Data

**DOI:** 10.1101/2021.03.31.437901

**Authors:** Doaa Hassan, Daniel Acevedo, Swapna Vidhur Daulatabad, Quoseena Mir, Sarath Chandra Janga

## Abstract

Pseudouridine is one of the most abundant RNA modifications, occurring when uridines are catalyzed by Pseudouridine synthase proteins. It plays an important role in many biological processes and also has an importance in drug development. Recently, the single-molecule sequencing techniques such as the direct RNA sequencing platform offered by Oxford Nanopore technologies enable direct detection of RNA modifications on the molecule that is being sequenced, but to our knowledge this technology has not been used to identify RNA Pseudouridine sites. To this end, in this paper, we address this limitation by introducing a tool called Penguin that integrates several developed machine learning (ML) models (i.e., predictors) to identify RNA Pseudouridine sites in Nanopore direct RNA sequencing reads. Penguin extracts a set of features from the raw signal measured by the Oxford Nanopore and the corresponding basecalled k-mer. Those features are used to train the predictors included in Penguin, which in turn, is able to predict whether the signal is modified by the presence of Pseudouridine sites. We have included various predictors in Penguin including Support vector machine (SVM), Random Forest (RF), and Neural network (NN). The results on the two benchmark data sets show that Penguin is able to identify Pseudouridine sites with a high accuracy of 93.38% and 92.61% using SVM in random split testing and independent validation testing respectively. Thus, Penguin outperforms the existing Pseudouridine predictors in the literature that achieved an accuracy of 76.0 at most with an independent validation testing. A GitHub of the tool is accessible at https://github.com/Janga-Lab/Penguin.

**HIGHLIGHTS:**

- Penguin integrates several developed ML learning models (i.e., predictors) to identify RNA Ψ sites in Nanopore direct RNA sequencing reads.
- The pipeline of penguin automates the data preprocessing including Nanopore direct RNA reads alignment using Minimap2, and Nanopore signal extraction using Nanopolish, feature extraction from raw Nanopore signal for training ML predictors integrated in its platform, and the prediction of RNA Ψ sites with those predictors.
- Penguin can predict Ψ sites with a performance that outperforms the performance of the state-of-the-art research methods existing in the literature.
- Penguin platform can be adopted to be used for predicting other/various types of RNA modification.
- There are 6137606 U-mers samples predicted by penguin best ML model (SVM) as Ψ ones from a total of 67491289 U-mers samples in the complete Hek293 cell line with 556813 unique genomic location of Ψ.
- There are 1193192 U-mers samples predicted by penguin best ML model (SVM) as Ψ ones from a total of 229637931 U-mers samples in the complete Hela cell line with 39384 unique genomic locations of Ψ.
- There is a small fraction of 0.01% (6482 unique genomic locations) of Ψ that are common (overlapped) between both Hek293 and Hela cell lines.
- The extend of Ψ modification (the number of U-mers samples predicted as Ψ samples to the total number of U-mer samples in the complete RNA sequence of the cell line) in RNA sequence of Hek293 cell line is much greater than its counterpart for Hela cell line (9% for Hek293 versus 0. 5 % for Hela cell line).

## 1. Introduction

Pseudouridine (abbreviated by the Greek letter Ψ) is one of the most abundant RNA modifications, occurring when uridines are catalyzed by Pseudouridine synthase proteins. It plays an important role in many biological processes such as stabilizing RNA through enhancing the function of transfer RNA and ribosomal RNA [1]. The modification of uridine to Ψ has been observed in nearly all kinds of RNA, including but not limited to tRNA (transfer RNA), and mRNA (messenger RNA). Ψ has also important roles in drug development and response to stresses [2].

Despite improvements in experimental technologies, chemical methods developed to detect Ψ sites, they remain both time-consuming and expensive. For example Ψ can be mapped by chemical conversion with N-cyclohexyl-N-(2-morpholinoethyl) carbodiimide metho-p-toluenesulphonate (CMC), which then blocks reverse transcription (Ψ-seq or pseudo-seq [1,3]). Ψ-seq detected 89 and 353 modified mRNA transcripts in different human cell lines. However, the number of sites identified with Ψ-seq is low compared with the spread of Ψ reported by mass spectrometry (MS) Methods, and the overlap between the different studies is very modest. Moreover, a method employing a chemical pulldown enrichment step for Ψ (N3-CMC-enriched Pseudouridine sequencing, CeU-seq) recognized 1929 modified mRNAs, but whether most of these sites were missed by Ψ-seq or are false positives has not been yet determined [4]. Therefore, there was a direction in the literature to address this problem using some computational biology methods that rely on developing machine learning algorithms to identify Ψ sites [5–11]. However, the performance results of such methods are quite low and they can be improved more.

Recently, the third-generation sequencing technologies such as the platforms provided by Oxford Nanopore Technologies (ONT) has been proposed as a new mean to detect RNA modifications on long read RNA sequence data. However this technology has not been used to identify Ψ sites in RNA sequence. To this end, our work aims to address this with a tool called Penguin. Penguin integrates several developed ML models (predictors) to identify Ψ sites in Nanopore direct RNA sequencing reads.

Penguin extracts a set of features from the raw signal of Oxford Nanopore RNA Sequencing reads and the corresponding basecalled k-mers. Those features are used to train the predictors included in Penguin, which in turn, is able to predict whether the signal is modified by the presence of Ψ sites. The features extracted by Penguin include: the signal length, some signal statistical features including the mean and standard deviation of the signal, and one-hot encoding of reference k-mers produced from aligning nanopore events/signals to a reference genome using the eventalign module in Nanpolish, a software for Nanopore signal analysis [12].

The developed predictors of Penguin have been trained and tested upon a set of ‘modified’ and ‘unmodified’ sequences containing Ψ at known sites or uridine respectively. All the predictors can be adopted to detect other RNA modifications which have not been yet tested.

Current trained predictors of Penguin are Support Vector Machine (SVM), Random Forest (RF), and Neural Network (NN). All the predictors have been trained and tested on two benchmark datasets for two human cell lines (namely the Hela and Hek293). However, Penguin can be also trained and tested with any other cell lines.

## 2. Materials and Methods

### 2.1 The pipeline of Penguin

The complete pipeline of Penguin is composed of various components/blocks (Figure 1). Penguin takes the fastq reads file that is generated from basecalling the fast5 files using any basecalling software (e.g., guppy, albacore, scrappy), and a reference genome as inputs. The fast5 files are produced by ONT device (these files are used to store the output of Nanopore sequencers and contains the raw electrical signal levels that come off the sequencers). Using both inputs to Penguin, the SAM file is created from the alignment stage (where the fastq reads file is aligned to reference genome to create this file). Next, using the produced SAM file and a provided BED file [13] (a file that highlights the target modified locations from the whole genome), the tool can create a coordinate file with ids of fast5 files that have the target modification. Next, the tool launches the Nanopolish software that performs signal extraction in two steps in order to produce a dataset of Nanopore signal samples. The first step takes the fast5 files folder as an input and index all files in that folder. The second step, takes the fastq reads file of the indexed fasts5 files, the sorted bam file (a sorted compressed version of the SAM file that is created using samtools software [14]), and the reference genome as inputs and run Nanopolish eventalign module on these inputs to generate a dataset of Nanopore signals.

**Figure 1.**
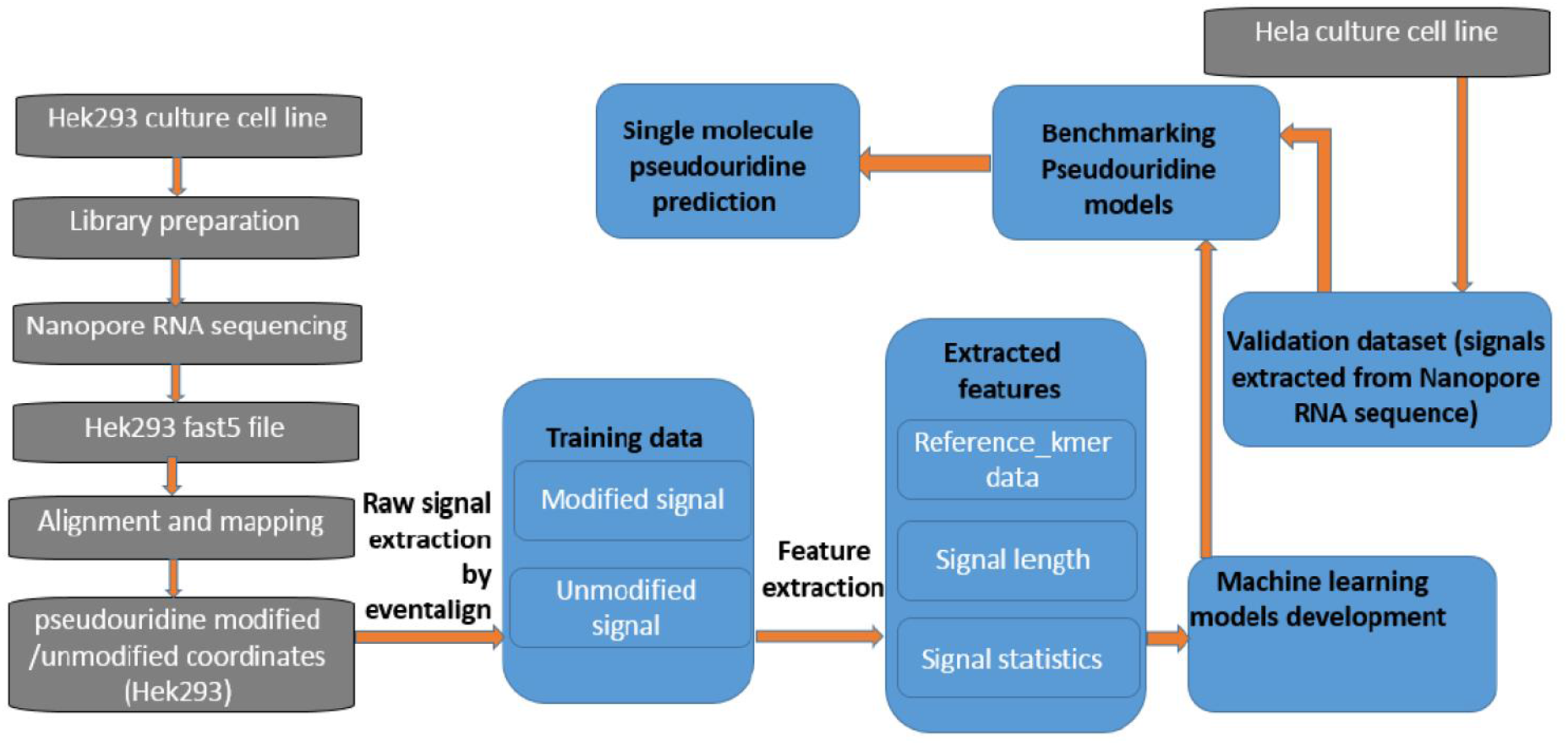
The complete pipeline of Penguin showing: the RNA sequencing of a HEk293 cell line, basecalling-alignment and mapping, identifying modified and modified Ψ coordinates, the raw signal extraction by nanopolish eventalign module, the feature extraction from signals that address Ψ modification, machine learning model development and validation, and single molecule Ψ prediction.

Next the signal samples that are related to Ψ modification are filtered from the samples generated from signal extraction. Using the information in the coordinate file some of the filtered samples will be labeled as modified and the remaining will be labeled as unmodified, as will be described later. Next some features will be extracted from those modified and unmodified samples which are fed to one of Penguin’s machine learning classifiers in the training phase. Using those features the classifier will be able to predict the modification sites in the testing phase. The details of each Penguin component will be described in the next subsections.

### 2.2 Cell Culture

Culturing the cells is where we extract cells from an animal and let it grow in an artificial environment [15]. Cell culture Hek293 and HeLa cell line were purchased from ATCC cell line collection, cultured in DMEM media supplemented with 10% FBS and 0.5% penicillin/streptomycin and grown at 37°C for 24-48hrs.

### 2.3 RNA library preparation and Nanopore RNA sequencing

The libraries that we used were prepared following Nanopore Direct RNA sequencing kit documented protocol (SQK-RNA002). Briefly, total RNA was isolated using Qiagen RNeasy Mini Kit (Cat No. /ID: 74104), followed by PolyA enrichment using Thermo Fisher Dynabeads™ mRNA DIRECT™ Micro Purification Kit (61021). 500 ng of poly (A) RNA was ligated to a poly (T) adaptor using T4 DNA ligase. Following adaptor ligation, the products were purified using Mag-Bind^®^ TotalPure NGS Beads (M1378-00), following NGS bead purification protocol. Sequencing adaptors preloaded with motor protein were then ligated onto the overhang of the previous adaptor using T4 DNA ligase followed by NGS bead purification protocol. The RNA library was eluted from the beads in 21 μl of elution buffer and quantified using a Qubit fluorometer using the manufacturer’s RNA assay. The final RNA libraries were loaded to FLO-MIN106 flow cells andrun on MinION. Sequencing runs and base calling was performed using MinKNOW software (Oxford Nanopore Technologies Ltd.), a software that controls the MinION Nanopore sequencing device. Therefore, once we extract the RNA from the cells during library preparation, we put it through the MinION device and start generating signal data. Nanopore is a small pore in MinION device that is used to create signal on biological molecules that pass through it. The MinION is one of the nanopore sequencing devices and it can read both DNA and RNA signals. When a strand of DNA or RNA goes through the pore it disrupts a membrane and creates a current that is measured by the device [16]. The output of the device is a fast5 file or files. These files contain the raw signals of the DNA or RNA reads.

The data output from MinKNOW for Hela cell line was 800k sequence reads using one flow cell. MinKnow generates data as pass and fail folders. FAST5 files from pass folder were again base called using Albacore 2.1.0 (Oxford Nanopore Technologies Ltd.) resulting in 723kb single molecule reads for Hek293 and 500K single molecule reads for Hela cell line which corresponded to full length transcripts ranging from 50b to 8kb. The directRNA-sequencing data generated for Hek293 and Hela cell lines in this study is publicly available on SRA, under the project accession PRJNA685783 and PRJNA604314 respectively.

### 2.5 Basecalling, alignment and mapping

There are several basecallers such as scrappy, albacore, guppy and many more [17]. The basecaller uses algorithms to determine what base is being passed through the Nanopore. Some fast5 files may or may not contain sequence information, so the basecallers can provide that for us. Once we have the sequence information, we can align the sequence to a reference genome. Aligning a sequence is just mapping the sequence to a certain location in the genome. It could be in any of the chromosomes and this information is saved to the SAM file.

### 2.6 Identifying Pseudouridine modified /unmodified coordinates

For a gold standard set of modified locations we use a BED file that address Ψ RNA modification locations in human genome (as Penguin has been tested on RNA data of two human cell lines: Hela and Hek239). These locations have been mentioned and verified in literature. The aforementioned BED file was imported from Epitomy, a gold standard RNA modification database that is used to search for locations of modifications [18]. Epitomy database incorporates data from over 51 different sources for human and mouse genomes at single nucleotide resolution and features more than 3 million modification loci in more than 12 RNA modification types. So Penguin uses these Ψ RNA modified locations imported from Eoitomy in a form of BED file to verify its prediction accuracy of Ψ sites. The imported Ψ BED file used Penguin has 2987 unique locations distributed in 25 unique genes (Supplementary text1).

### 2.7 Raw signal extraction

The Nanopore direct RNA sequencing holds a great advantage in identifying RNA modifications at single base resolution by the interpretation of ONT raw signals ( i.e., events or “squiggles”) plotted over time that are corresponding to modified and unmodified base sequence contexts. To this end, the basic functionality of Penguin aims to exploit the difference in the raw signal between modified and unmodified bases in order to predict Ψ sites in RNA sequence. Thus, the signal extraction phase plays an important role in achieving Penguin functionality. For this phase, we use Nanopolish [19], the ONT analysis software. In particular, the eventalign module [20] of Nanoploish is used to extract the raw Nanopore signals and get some of its corresponding features such as the mean, standard deviation and length of the signal. In order to achieve this, eventalign first aligns events to a reference genome, and so the low-level signal information can be obtained. Such information can be used to discover the differences in the current that might lead to base modifications. We refer to Nanopolish manual for more information about the Nanopolish pipeline for signal analysis [21].

### 2.8 Benchmark datasets generation

Two different benchmark datasets were generated for Hek293 and HeLa cell lines (Supplementary csv files Hek293.csv and Hela.csv). In order to generate those datasets, we consider the samples of eventalign output (where eventalign was run on the basecalled sequence reads of each cell line) as the samples of the benchmark. In order to label each sample, we filter all samples that have a U base in the middle of the model k-mer (one column in event align output), which is the target base for identifying Ψ modification. Next, we find the intersection between their position column on the reference genome and the position in the coordinate file (generated from Ψ BED file and SAM file for each cell line). This intersection will represent the positive samples, while the remaining samples will be the negative samples. In the end we have 13072 samples: 6536 are positive and 6536 are negative ones (after sampling the negative samples which are very huge in comparison with negative ones) for Hek293. Similarly, we get 1354 samples: 677 are positive samples and 677 are negative samples for HeLa cell line.

### 2.9 Feature extraction

Each generated benchmark dataset has 4 columns that represent four features that were used for training the machine learning models that we developed and integrated in Penguin platform. Those features were extracted by picking 4 columns from the evenalign output (Supplementary text2) (namely: reference_kmer, mean, stdv, and length of event) and use them as features for training the constructed ML classifiers of Penguin. Those columns refer to the k-mers generated from aligning events to a reference genome, the mean, the standard deviation, and the length of each extracted signal respectively.

### 2.10 ML Models construction

We have developed several machine learning models and integrate them into Penguin’s platform including the SVM [22], RF [23] and NN [24]. Those algorithms have been used extensively to address several problems in bioinformatics research [25], [26], [27]. The radial basis function kernel (rbf) was used in SVM training. The gamma parameter was set to ‘scale’ and the default value of C parameter was used. For RF, the seed number was set to 1234 and the number of trees was set to 30. As for NN, a two hidden layers NN was implemented. The number of neurons (nodes) was 12 in the first and 8 in the second. The ‘Adam’ optimizer with a learning rate of 0.001 was used, the number of epochs were set to 150, and the batch size was set to the length of training set.

We have used the scikit-learn toolkit [28], the free machine learning python library to implement the SVM and RF models, while a keras [29] with tensorflow [30, 31] back end were used to implement the NN model.

### 2.11 Feature importance

As for deep analysis of the features that contribute mostly to the performance of machine learning model that does Ψ site identification, we have reported about the importance of each feature thorugh training and testing each developed ML model of Penguin with each of the four extracted features that were described in sec 2.9. This is acheived by buliding five versions of the ML model applied to each benchmark dataset. In the first four versions, each developed ML model is trained with one of the four extracted features, while the in fifith version each model is trained with the combination of all four features.

### 2.12 Random test splitting

In this approach, we randomly divide the benchmark datasets into two folds: one is for training and another one for testing. The test_size was set to 0.2 which means 80% of the benchmark dataset is used for training the model and 20% of the dataset is kept for testing the model.

### 2.13 Test with independent cell line

In this approach two benchmark datasets for two different cell lines are used one is used for training and another one is used for testing.

### 2.14 Evaluation metric

We have used the accuracy (Acc), precision, recall, and the area under the curve (AUC) [32] as metrics to evaluate the performance of penguin predictor. Below we introduce the mathematical equation for the first three metrics:

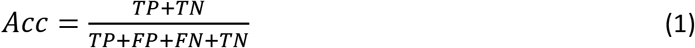

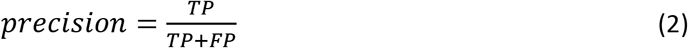

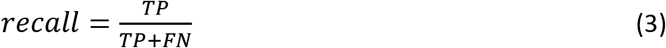

Where:

- TP stands for true positive and refers to the number of correctly classifed Pseudouridine sites.
- FP stands for false positive and refers to to the number of non Pseudouridine sites that were misclassified as Pseudouridine sites.
- FN stands for false negative and refers to the number of Pseudouridine sites that were misclassified as non Pseudouridine sites
- TN stands for true negative and refers to the number of correctly classifed non Pseudouridine sites.

## 3. Results

We have used the two validation methods: the random-test splitting and the test with independent cell line introduced in Section 2 for evaluating the performance of each predictor in Penguin platform using the metrics that we previously mentioned. In the following we present the performance results that we have got using each method.

### 3.1 Performance evaluation with random-test splitting

Table. 1 shows the performance of ML models (Supplementary python files SVM.py, RF.py, and NN.py) using the extracted 4 features introduced in Section 2 from the benchmark dataset of Hek293 cell line. Clearly SVM achieves the best performance in terms of accuracy as it achieves 93.38%, while RF achieve best precision of 0.98 and SVM and NN achieve the best recall of 0.95 per each.

**Table 1:**
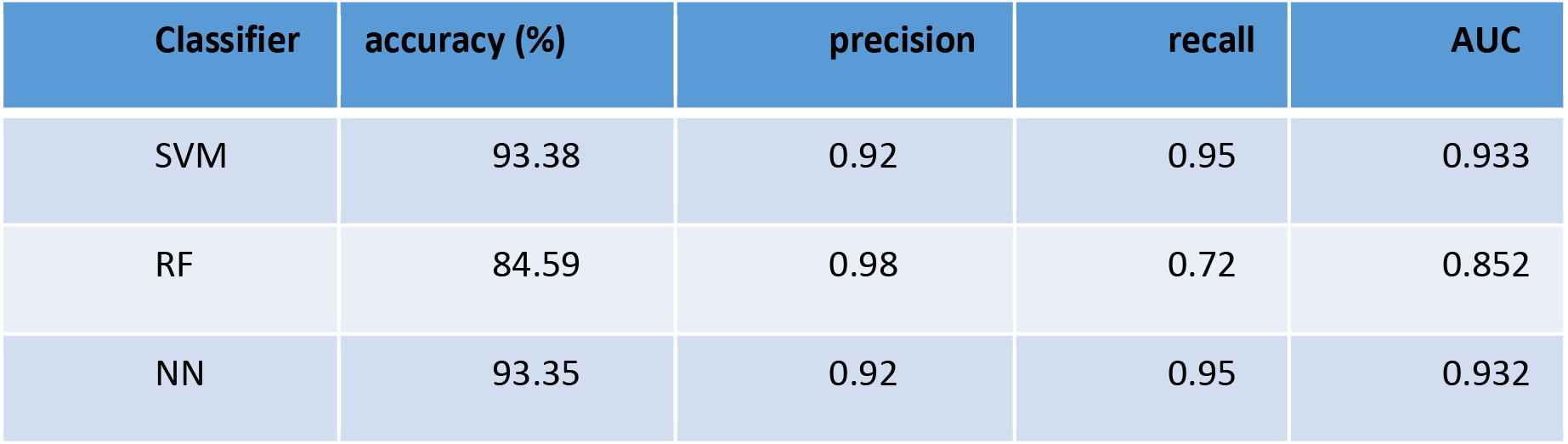
The performance of Penguin’s predictors on Hek293 benchmark dataset with random-test splitting using all four extracted features.

The learning curve of SVM, RF, and NN (see Figure 2, panels A,B, and C respectively) show the performance of SVM and NN in terms of accuracy score outperforms the perofrmnce of RF. Also we used receiver operating characteristic (ROC) which plots the true positive rate on the vertical coordinate versus the false positive rate on the horizontal coordinate. The ROC curve of SVM, RF and NN (see Figure 3 A,B ansd C) show that the percentage of true postive rate to the fasle positive rate in case of SVM and NN is more that the case of RF.

**Figure 2.**
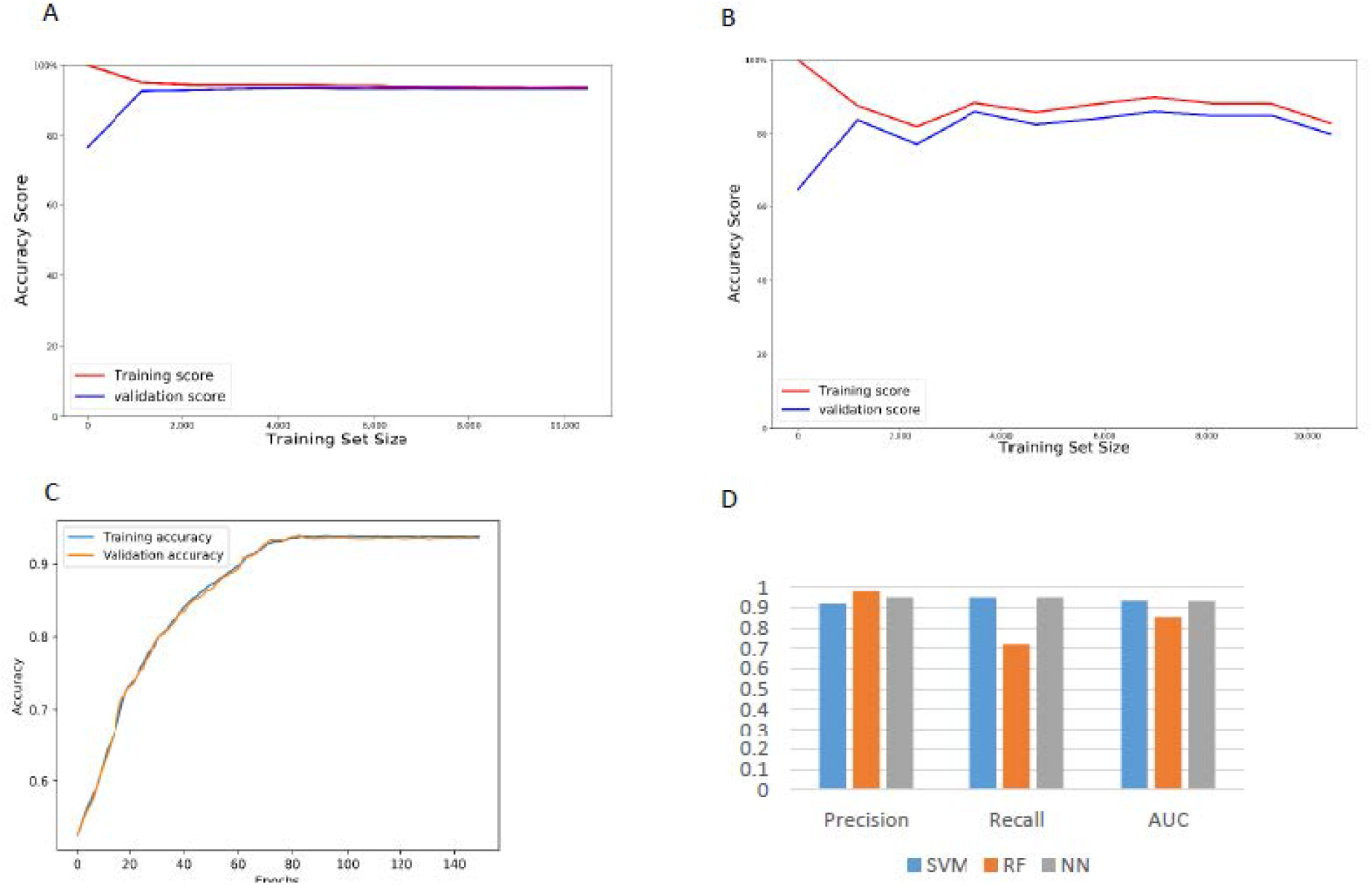
The learning curve for each Penguin’s predictor and a bar chart of the performance evaluation using Hek293 cell line benchmark dataset with random test splitting showing (a) The learning curve of SVM (b) The learning curve of RF (c) The learning curve of NN (d) A bar chart of the precision, recall, and AUC of Penguin’s developed ML classifiers.

**Figure 3.**
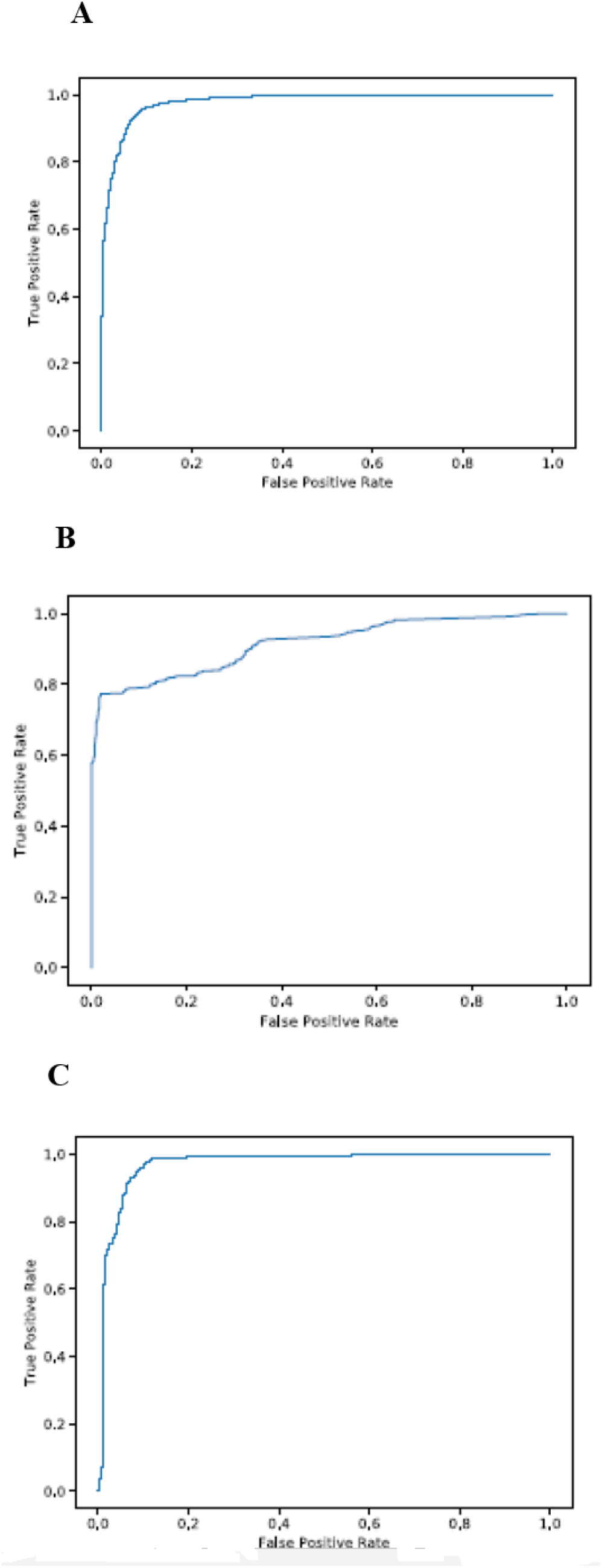
The ROC curve for each Penguin’s predictor using Hek293 cell line benchmark dataset showing (a) The ROC curve of SVM (b) The ROC curve of RF (C) The ROC curve of NN.

#### 3.1.1 Performance results using single type of feature

Table 2 shows the performance of ML models with random test-splitting on Hek293 benchmark dataset in terms of accuracy with single type of feature among the four extracted features introduced in Section 2.

**Table 2:** The performance of Penguin’s predictors on Hek293 benchmark dataaset in terms of accuracy with random test-splitting using single type of feature.

| Classifier | Mean | Stdv | Length | One-hot Encoding | Combination |
| --- | --- | --- | --- | --- | --- |
| SVM | 69.33 | 61.87 | 51.81 | 93.38 | 93.38 |
| RF | 69.17 | 61.49 | 50.86 | 87.92 | 84.59 |
| NN | 67.34 | 46.46 | 47.53 | 93.42 | 93.35 |

Clearly the one-hot encoding of reference_kmer contributes more to the classifier’s accuracy than other features for all classifiers. It is by followed by mean, then standard deviation, then the length feature comes at the end for SVM and RF and the standard deviation comes at the end for NN. Based on the results in Table 2, one would wonder why we do not use the one-hot encoding feature alone for prediction and avoid using features extracted from the signal as the former leads to a classification accuracy equal to that when using the combination of four features in case of SVM and outperforms the classification accuracy of combination in case of RF and NN. However, using onehot encoding feature alone is not useful when predicting new location of Ψ sites that have not seen before. Also the extracted signals are still needed even if their features are not contributing to the performance as one-hot encoding feature. This is because without signals extraction we can’t get the model_kmer that is needed to filter the samples in eventalign output that address the target modification.

### 3.2 Performance against independent cell line

Table 3 shows the performance of ML models against independent cell line (i.e., with independent test dataset, where Hek293 cell line benchmark dataset is used for training penguin’s predictors and Hela cell line benchmark dataset is used for testing them) using the four extracted features introduced in Section 2. We only include SVM and NN models (Supplementary python files SVM_validate.py and NN_validate.py) and execlude the RF model as it acheives very low performance against independent cell line (below 50% accuracy) in comparison with SVM and NN. Clearly as shown in Table3 NN outperforms SVM. However, SVM is more stable as it achieves reproducable results, while NN performance results changes from run to run though all NN results are still high (above 93% accuracy). See also learning curves of SVM and NN (Supplementary Figure 1 and Figure 2 respectively) and ROC curves of SVM and NN (Supplementary Figure 3 and Figure 4 respectively).

**Table 3:**
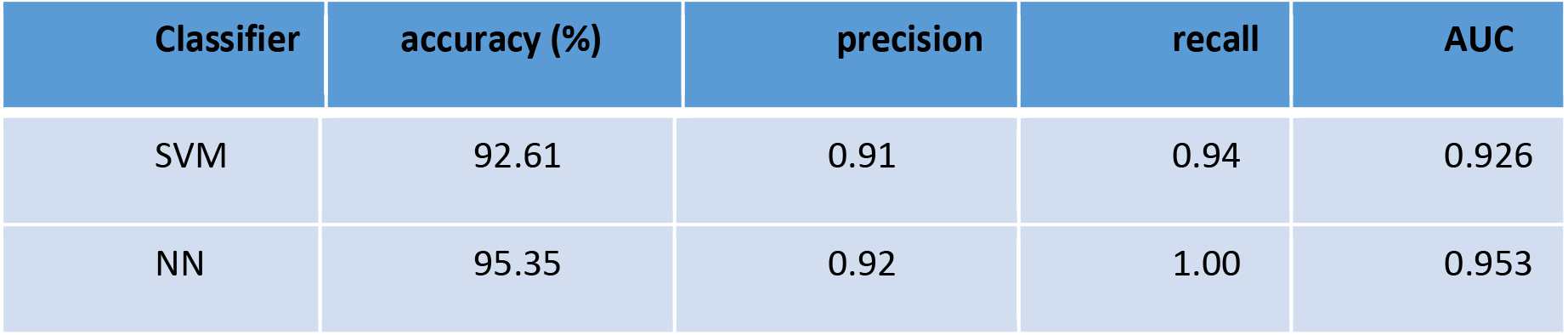
The performance of Penguin’s predictors against independent cell line using all extracted features.

| Classifier | accuracy (%) | precision | recall | AUC |
| --- | --- | --- | --- | --- |
| SVM | 92.61 | 0.91 | 0.94 | 0.926 |
| NN | 95.35 | 0.92 | 1.00 | 0.953 |

#### 3.2.1 Performance results using single type of feature

Table 4 shows the performance of ML models against independent cell line in terms of accuracy with single type of feature among the four extracted features introduced in Section 2. Clearly the one-hot encoding of reference_kmer contributes more to the classifier accuracy than other features, followed by length, then standard deviation, then mean feature comes at the end.

**Table 4:** The performance of Penguin’s predictors with against independent cell line using single type of extracted features.

| Classifier | Mean | Stdv | Length | One-hot Encoding | Combination |
| --- | --- | --- | --- | --- | --- |
| SVM | 39.22 | 40.99 | 51.62 | 95.35 | 92.61 |
| NN | 32.94 | 37.59 | 51.18 | 95.35 | 95.42 |

## 4. Abundance of Ψ sites

In order to identify the abundance of Ψ sites in the RNA sequence of either Hek293 or HeLa cell lines, first we run the best Penguin’s machine learning model (i.e., SVM as it achieves very high reproducible performance results over all runs) on the complete RNA sequence reads of Hek293 and HeLa cell lines. Then, we identify all U-mers samples predicted as Ψ ones in those reads, then we identify the number of Ψ unique genomic locations as well as their frequencies in the two complete cell lines. We found that there are 6137606 U-mers samples predicted as Ψ ones from a total of 67491289 U-mers samples of the complete RNA sequence of Hek293 cell line with 556813 unique genomic location of Ψ (Supplementary excel file 1)^1^. Similarly, we found that there are 1193192 U-mers samples predicted as Ψ ones from a total of 229637931 U-mers samples in the complete RNA sequence of Hela cell line with 39384 unique genomic locations of Ψ (Supplementary excel file 2). As for overlapping between unique genomic locations of Ψ in both cell lines, we found 6482 unique genomic locations of Ψ that are common between both cell lines (Figures 4.A). Similarly, we found that there are 7148 modified Ψ genes that are common/overlapped between both cell lines. Also, we found that there is an overlapping of 15.8% between the top 1% frequent modified Ψ genes of both complete Hela and Hek293 cell lines (Figures 4.B).

**Figure 4.**
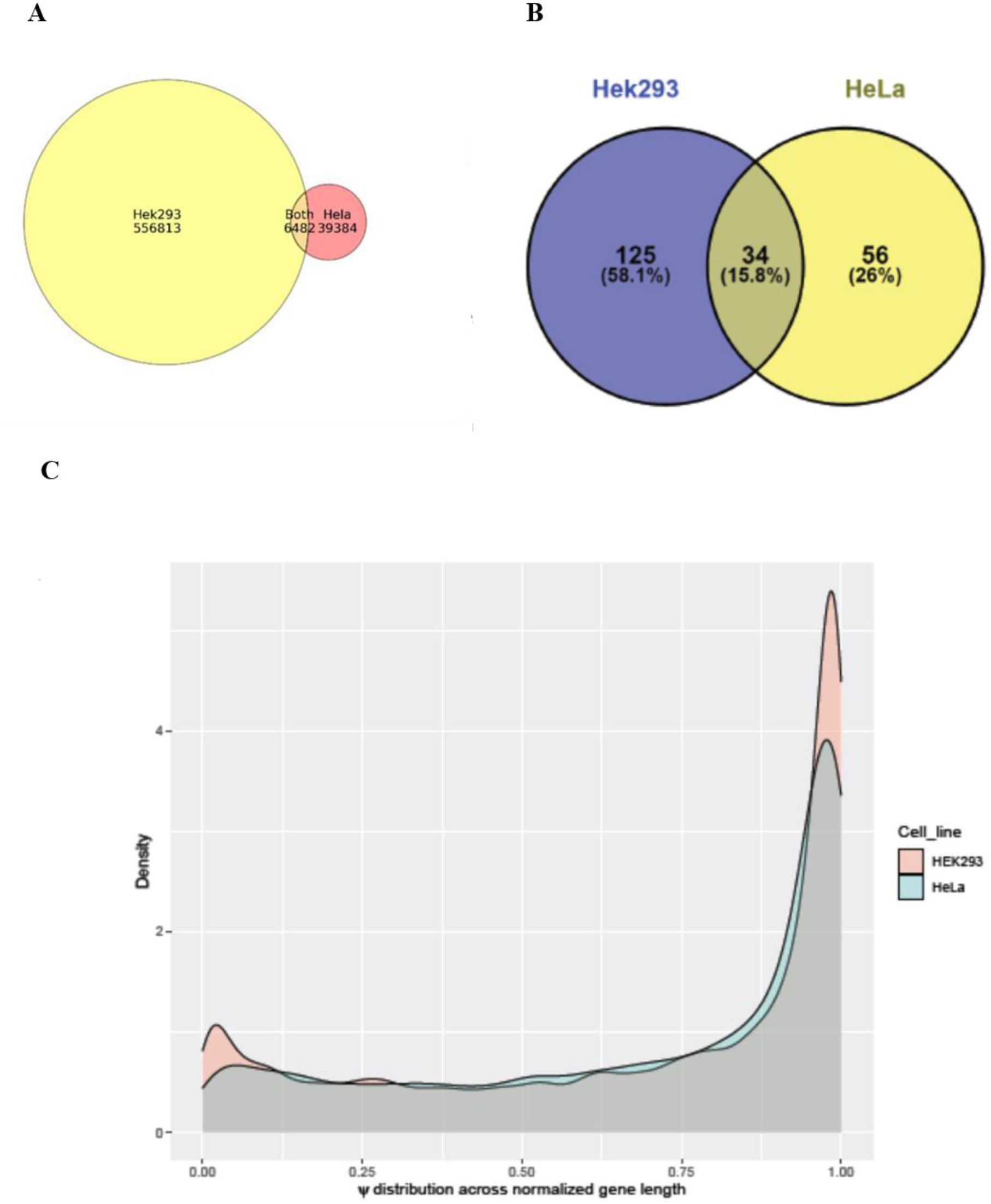
Showing (a) The overlapping between *Ψ* unique locations in complete Hek293 and HeLa cell lines (b) the overlapping between top frequent 1 % modified Ψ genes in complete Hek293 and Hela cell lines (c) The density plots that represents ψ distribution across normalized gene length for Hek293 and Hela cell lines.

Clearly, we notice that the extent of Ψ modification (the number of U-mers samples predicted as Ψ samples to the total number of U-mer samples in the complete RNA sequence of the cell line) in RNA sequences of Hek293 cell line is much greater than its counterpart for HeLa cell line (9% for Hek293 versus 0. 5 % for Hela cell line). This is due to the existence of more modified Ψ genes with extensive Ψ unique genomic locations in RNA sequences of the complete Hek293 as we just mentioned in comparision to the number of modified Ψ genes with Ψ fewer unique genomic locations found in the RNA sequence of the complete Hela cell line. Therefore the ψ distribution across normalized gene length for Hek293 cell line is larger than it equivalent in HeLa cell line (Figures 4.C).

## 5 Functional enrichment analysis

To investigate the potential functional role of Ψ modification in RNA, we performed functional enrichment analysis for the most frequently modified Ψ genes (top 1%) across Hek293 and HeLa cell lines. A total of 159 genes from Hek293 and 90 genes from Hela cell lines were identified to have the most abundance of Ψ RNA modification. The short-listed genes from both cell lines were plugged into Cytsoscape ClueGo [33] application to obtain enriched ontologies and pathways at high confidence (p<0.05). Enrichment observations from this analysis are visualized in Figure 5.

**Figure 5.**
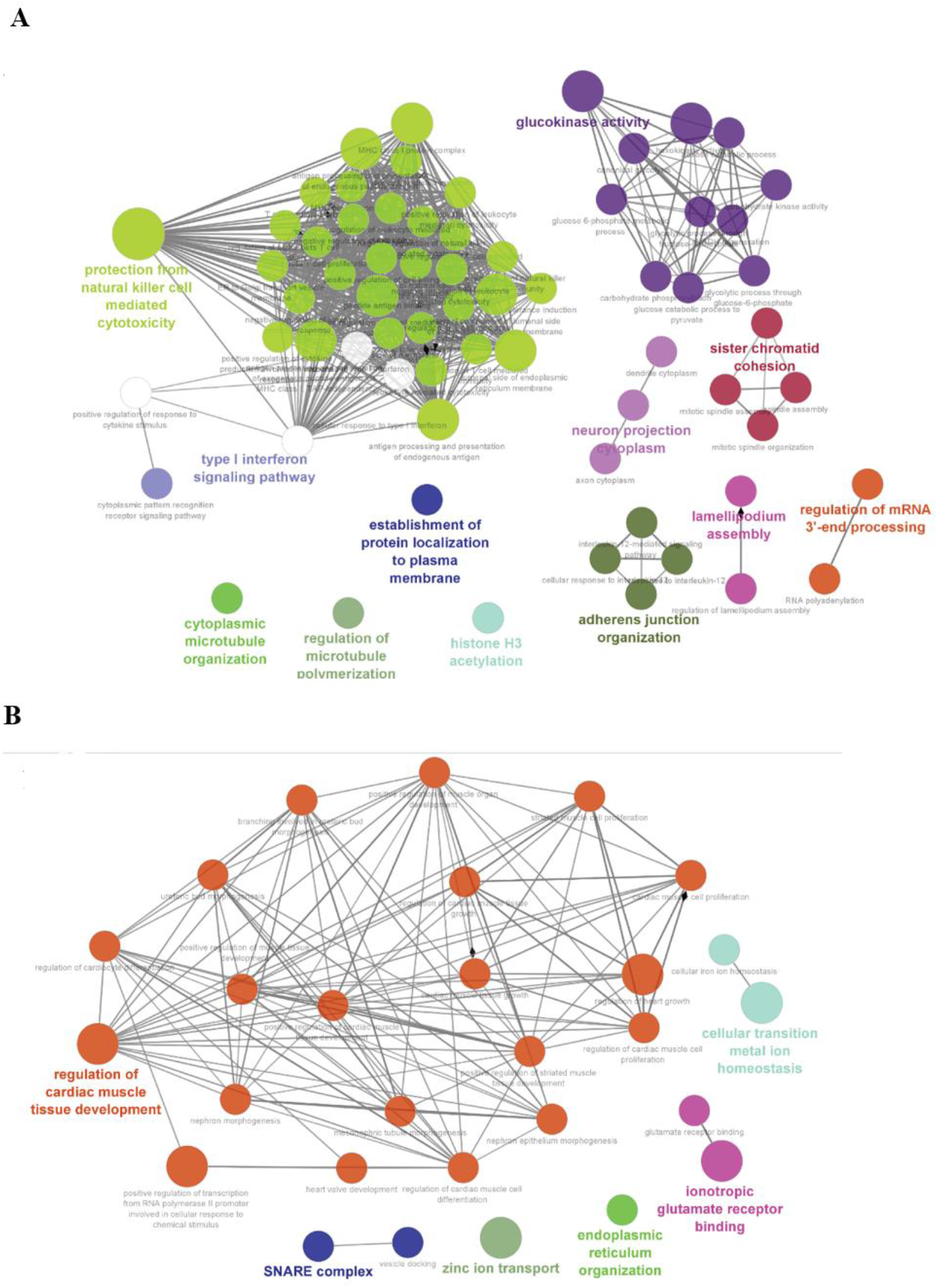
Functional enrichement analysis of most frequenlty Ψ modified genes across (a) Hek293 cell line and (b) Hela cell line, visualizing high confidence (p-val<0.05) ontologies and pathways potentially associated with Ψ RNA modification. Cliques clustered by functional grouping of the GO-terms based on GO hierarchy using Cytoscape ClueGO application. Size of the nodes representative of the significance of association with respect to genes per GO-term.

From the functional enrichment analysis of the gene set form Hek293 cell line, we observed a wide range of functional processes like (as seen in Figure 5B): “protection from natural killer cell mediated cytotoxicity”, “T cell mediated cytotoxicity”, “glucokinase activity”, “histone H3 acetylation”, “type I interferon signaling pathway”, and “regulation of mRNA 3′-end processing” being significantly (adjusted p-val < 0.0013) enriched. Essentially highlighting the diverse regulatory role of Ψ modification, from its involvement in cell immune signaling to mRNA 3’-end processing.

In HeLa cell line, we observed several high confidence (adjusted p-val < 0.0014) enriched ontologies that were more representative of Ψ modification role in cellular development and homeostasis via post-transcriptional regulation (as seen in Figure 5B): “regulation of cardiac muscle tissue development”, “cellular transition metal ion homeostasis”, “positive regulation of transcription from RNA polymerase II promoter involved in cellular response to chemical stimulus”, “SNARE complex”, “Zinc ion transport”, and “endoplasmic reticulum organization”.

## 6. Discussion

Penguin tool main file is implemented in python 3.x and the tool has to be run on Linux environment by running the following command from Penguin main directory on the user’s local machine after cloning the code from Penguin GitHub repository:

python main.py -r ref.fa -f reads.fastq

Where the penguin tool needs the following two inputs files when running it:

- The absolute path to the reference Genome file (ref.fa)
- The absolute path to fastq reads file (reads.fastq)

Once the user runs the penguin tool, then the tool pipeline (Figure 1) that accepts the aforementioned inputs will start execution and the user will be asked to enter the bed file name with the absolute path and extension that is needed to generate the coordinate file that is needed for labeling the U-mers Nanopore signals samples as modified and unmodified ones. Next, the tool will extract the raw Nanopore signals from the input fast5 file(s) as well as extracting some of its corresponding features that are used later to train the three different machine learning models (SVM, RF, and NN) integrated in Penguin platform for predicting Ψ sites in direct Nanopore RNA sequence. However this raises a question about why we did not develop a deep learning model for this prediction task? So the answer is simply because our developed machine learning models for Penguin achieved high accuracy in predicting Ψ sites and their performances significantly outperform the performances of other existing models in the literature that were developed for the same purpose. However, Penguin platform can also be extended to involve deep learning models for predicting Ψ sites or any other type of RNA modification in Nanopore sequencing data and we leave this as a future work.

## 7. Conclusions

In this paper, we have proposed a new tool called Penguin that represents a complete pipeline for predicting Ψ sites in direct Nanopore RNA sequence of reads. It has been shown that our proposed tool outperforms the existing prediction models of Ψ in the literature. We expect that Penguin will become a useful tool for accurate identification of Ψ sites in RNA read sequence. Penguin platform can be adopted to be used for predicting other types of RNA modification.

For future work, we are looking forward to address the general limitiation of the method used to detect Ψ modifications using Nanoppore technology which is implmented in Penguin. Such a limitation occures because the prediction of RNA modification in ONT sequencing is further complicated due to the fact that each sequencing signal originates from a group (k-mer) rather than from a single nucleotide. Furthermore, the signal for each k-mer is variable, leading to a distribution of possible signals. So RNA modifications can result in a shift in the signal distribution for a given k-mer, and this shift can be used to predict the presence of modifications. However, these shifts can be relatively small, and the distribution of signals generated by modified k-mers largely overlaps with the distribution of unmodified k-mers which leads to mis-identifying some Ψ sites [3]. Moreover, the modification prediction platform of Penguin that employs machine-learning models depends on the quality of the training datasets generated by the automated pipeline of penguin for data preprocessing including basecalling, alignment using Minimap2, and signal alignment using Nanopolish.

## Author Contributions

DH, DA, and SCJ conceived and designed the study. DH implemented the Penguin tool pipeline and its machine learning predictors. DH extracted the benchmark datasets and evaluated the performance of Penguin predictors with the random test split and against independent cell line. DH and SVD performed results and functional enrichment analysis. QM performed the cell cultural, RNA library preparation and Nanopore RNA sequence. DH and QM read and approved the final manuscript.

## Acknowledgement

We thank Rajashekar Varma Kadumuri form whom we borrow the code that we slightly updated to generate the coordinate file for Hela and Hek293. We also thank Alexander Krohannon and Ratanond Koonchanok at IUPUI for giving valuable comments on this work and valuable discussions. This work is supported by the National Science Foundation (NSF) grant # 1940422 and #1908992 as well as the National Institute of General Medical Sciences of the National Institutes of Health under Award Number R01GM123314 (SCJ).

## Footnotes

1 U-mers samples are rows (with U in the middle of their reference kmers column) in the eventalign output that corresponds to the result of aligning the events (signals or squiggles) of RNA sequence of a specific cell line to a reference genome.

